# Hybrid Deep Learning with Protein Language Models and Dual-Path Architecture for Predicting IDP Functions

**DOI:** 10.1101/2025.05.25.655984

**Authors:** Jiahui Liang, Zhenling Peng

## Abstract

Intrinsically disordered regions (IDRs) drive essential cellular functions but resist conventional structural-function annotation due to their dynamic conformations. Current computational methods struggle with cross-dataset generalization and functional subtype discrimination. We present IDPFunNet, a hybrid deep learning model combining convolutional neural network, bidirectional LSTM, residual MLP, and the protein language model ProtT5 to predict six IDR functional classes: five binding subtypes and disorded flexible linkers (DFLs). Its dual-path architecture decouples binding IDR prediction and DFL identification using evolutionary embeddings from ProtT5, which outperformed ESM-family models and AlphaFold2 structural features by ≥ 1.3% average AUC and ≥ 12.7% average APS. Validated across six benchmarks including CAID2/3 blind tests, IDPFunNet achieves superior protein-binding prediction (AUC 0.832-0.866) with significant improvements (≥1.5%, p-value<0.05) over existing methods, while matching specialized DFL predictors. Multi-task learning enhances protein/lipid/small molecule-binding performance (3.1-35.1% AUC gains) with BiLSTMs proving optimal DFL identification, despite self-attention’s potential for nucleic acid-binding (AUC 0.831). This framework overcomes methodological limitations through interpretable, generalizable IDR functional mapping.

## Introduction

Intrinsically disordered proteins (IDPs) constitute a vital class of biomolecules that perform essential cellular functions through intrinsically disordered regions (IDRs), despite lacking stable tertiary folds under physiological conditions^1-6^. These conformationally plastic regions enable multifaceted interactions with diverse binding partners—from proteins to small molecules—through adaptive folding mechanisms ^7-9^. A prototypical example is the p53 transcriptional activation domain (TAD2), which binds diverse partners like S100β and CBP through conformational plasticity ^10^. Beyond direct molecular recognition, IDRs frequently serve as flexible linkers (DFLs) that mediate allosteric communication between structured domains ^11-13^. The p21 cyclin-dependent kinase inhibitor exemplifies this role, where its linker helix subdomain LH dynamically bridges D1 and D2 subdomains to accommodate structural variations across Cdk/cyclin complexes, thereby enabling precise cell cycle regulation ^14^.

The inherent structural plasticity of IDRs presents formidable challenges for high-throughput experimental characterization, necessitating computational approaches for systematic functional annotation. Early methodologies predominantly employed handcrafted features—including biophysical properties, sequence conservation patterns, and evolutionary coupling data—combined with classical machine learning algorithms (support vector machine, logistic regression, random forest) to predict IDR functions. Representative approaches from this era include ANCHOR ^15^, DisoRDPBind ^16^, MoRFchibi SYSTEM ^17^, DFLpred ^18^, APOD ^19^, IDPpi ^20^ and CLIP ^21^. This paradigm evolved with deep learning advancements, yielding hybrid frameworks like flDPnn^22,23^, DeepDISOBind ^24^, and DisoLipPred ^25^ that integrate neural architectures with traditional feature engineering. The critical assessment of intrinsic disorder (CAID) experiment ^26,27^ systematically benchmarks these methods, revealing intriguing evolutionary trends: while ensemble machine learning models like MoRFchibi SYSTEM ^17^ rivaled deep learning counterparts (e.g., DeepDRPbind ^28^) in CAID2 binding IDR prediction, deep neural networks dominated DFL prediction, with AlphaFold-based methods and SPOT-Disorder leading the rankings^29,30^. Recent breakthroughs in protein language models (PLMs), including ESM-family models ^31,32^ and ProtT5 ^33^ have revolutionized computational biology, demonstrating unprecedented capabilities across structure prediction ^32-37^, protein engineering ^38,39^, and variant effect analysis ^40,41^. PLM applications in disorder prediction include IDP-ELM’s ensemble strategy combining ESM and ProtT5 embeddings^42^, and DisoFLAG’s ProtT5-GNN hybrid achieving state-of-the-art (SOTA) performance across six functional categories ^43^. Despite these innovations, persistent challenges arise from dataset-specific annotation biases that compromise prediction consistency.

To address these limitations, we present IDPFunNet - a dual-path neural architecture that utilizes the hybrid of convolutional neural network (CNN) ^44^ and bidirectional long short-term memory (BiLSTM) ^45^ for binding IDR identification and the specialized BiLSTM model for DFL prediction. Our design rationale stems from the distinct biophysical signatures of these functional classes ^46-48^. Through systematic benchmarking against CAID2/3 leaders and ablation studies, we demonstrate IDPFunNet’s superior stability and performance in binding IDR prediction while matching specialized DFL predictors. We further quantify the differential impacts of AlphaFold2 ^49^ structural features, multi-task learning, and self-attention mechanisms ^50^, providing mechanistic insights into optimal model design for IDR multifunctional annotation.

## Results

### Overview of IDPFunNet

IDPFunNet is a hybrid deep learning framework, taking protein sequence as input, and integrating ProtT5 with dual-path architecture for multifunction prediction of IDRs. The framework encodes residue-level evolutionary features via ProtT5 embeddings, which are processed through two specialized paths: (1) a CNN-BiLSTM hybrid capturing local motifs and long-range dependencies for protein/nucleic acid/lipid/ion/other small molecule-binding IDR prediction, and (2) a BiLSTM network optimized for DFL identification. Feature vectors from both paths are fed into six parallel residual multilayer perceptron (ResMLP) blocks, generating probability scores *p* via sigmoid activation to quantify six functional propensities for each predicted residues. This architecture enables simultaneous prediction of binding interactions and structural linkers, addressing the heterogeneous nature of IDR functions through complementary feature extraction mechanisms.

### Insight on the CNN-BiLSTM complementarity for IDR multifunction prediction

CNN and BiLSTM architectures have become cornerstone architectures for protein function prediction ^51-55^. The CNN architecture specializes in local motif detection ^44^, while BiLSTM excels at modeling long-range sequence dependencies ^45^. To systematically evaluate their applicability to IDR multi-functions, we engineered standalone CNN and BiLSTM variants by substituting the original dual-path in our base model with their respective architectures (Figure 1). Performance was quantified via AUC and APS metrics on the TE210 benchmark.

**Figure 1.**
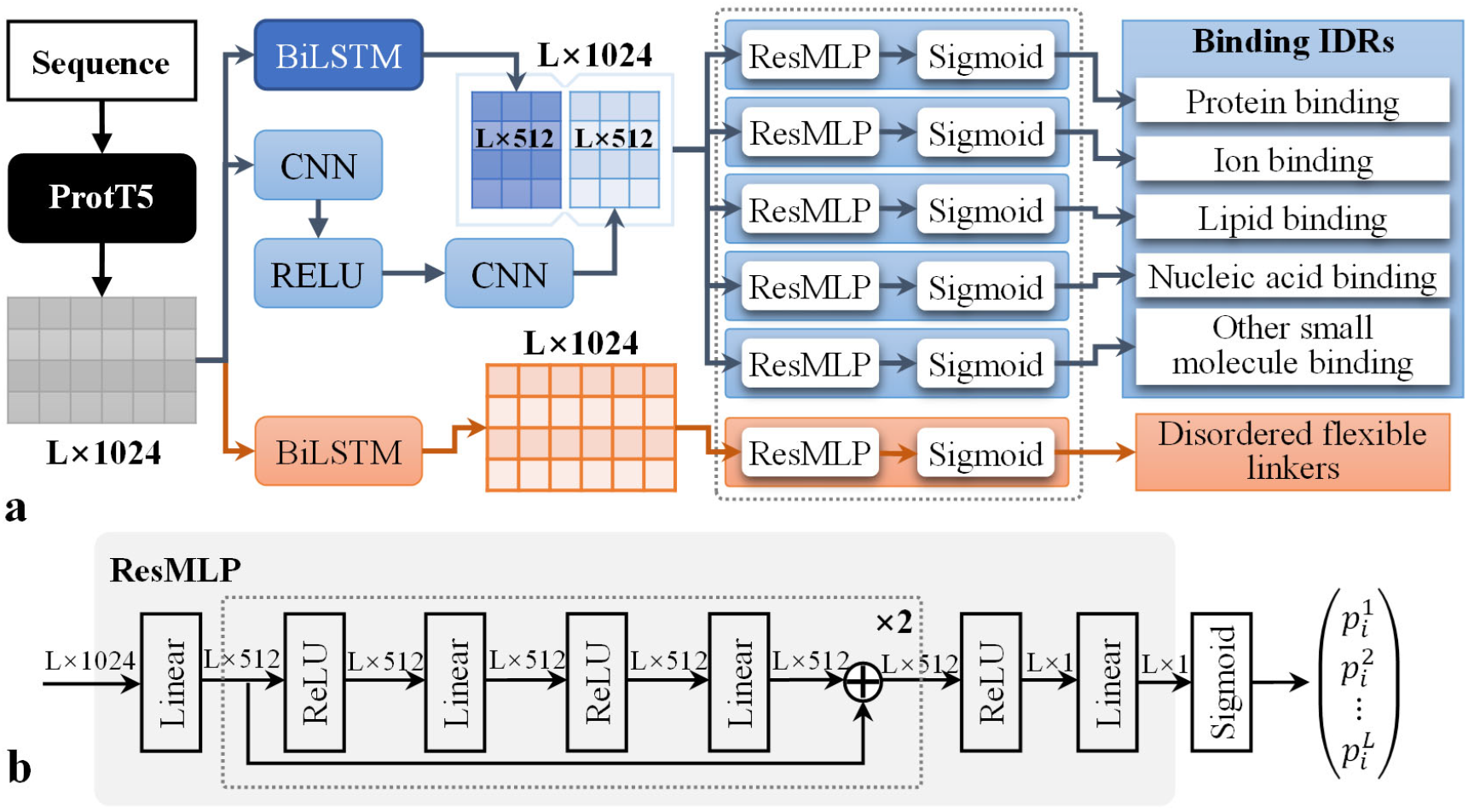
The architecture of the IDPFunNet model.

Comparative analysis (Figure 2) revealed both architectures achieved baseline performance (DFL prediction: AUC ∼0.88, APS 0.193–0.227; binding IDRs: AUC ∼0.78, APS ∼0.22). Crucially, we observed a functional dichotomy: BiLSTM outperformed in nucleic acid/lipid-binding prediction (0.3– 1.8% higher AUC and 25.3-75.9% higher APS), whereas CNN dominated protein/other small-molecule detection (0.8–4.1% higher AUC and 0.5–143.5% higher APS). This task-specific performance divergence directly correlates with their inherent strengths—local pattern recognition versus global context integration.

**Figure 2.**
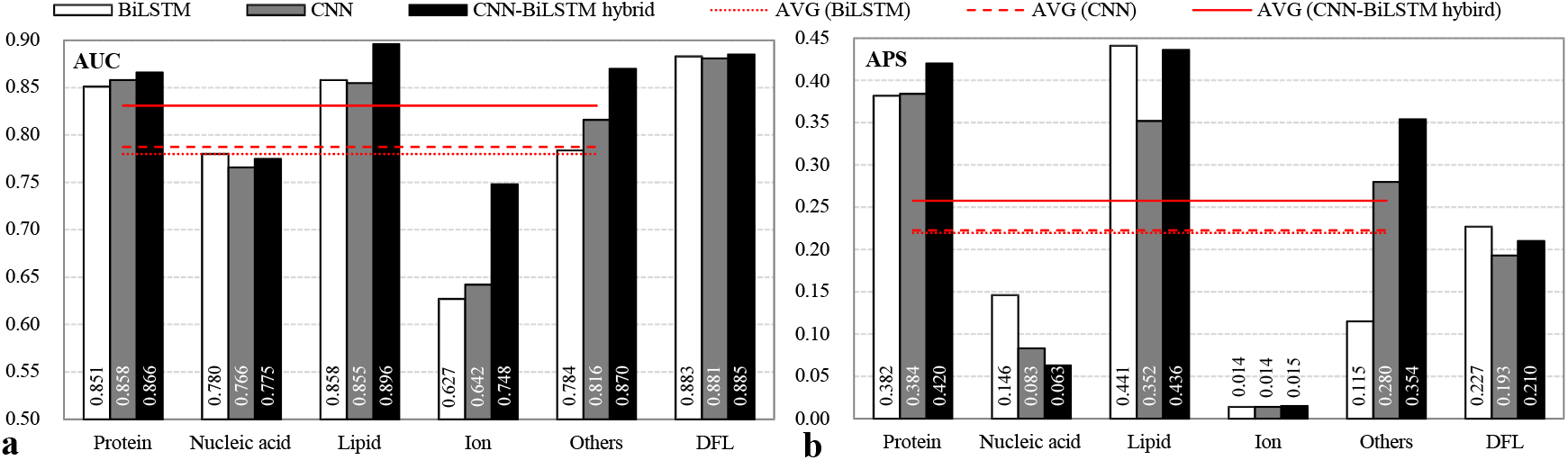
The predictive quality of the CNN-based, BiLSTM-based and the CNN-BiLSTM hybrid models, which is evaluated by using (**a**) AUC and (**b**) APS metrics on the test set TE210. These models were developed by replacing the original dual-path architecture in our base model IDPFunNet with their respective architectures. The “Others” category represents IDRs binding to other small molecules. The cross-category average AVG (*) aggregates performance across five binding classes: protein-, nucleic acid-, lipid-, ion-, and other small molecule-binding IDRs.

Hybridization of these architectures yielded significant improvements: The CNN-BiLSTM ensemble surpassed standalone models by 0.93–19.3% AUC across most binding tasks, with average AUC and APS improvements of ≥5.5% and ≥15.7%, respectively (Figure 2). Paradoxically, BiLSTM alone matched hybrid performance in DFL prediction (AUC ∼0.88) while achieving ≥8.1% higher APS. These findings demonstrate CNN’s complementary role to BiLSTM in binding IDR prediction but functional redundancy in DFL task.

Guided by the structural dichotomy between interaction-prone and linker IDRs ^27^, we formalized this specialization through IDPFunNet’s dual-path architecture: CNN-BiLSTM hybrids for binding IDR prediction and dedicated BiLSTM streams for DFL annotation.

### ProtT5 outperforms ESM-family models and AlphaFold2 in feature encoding

Recent advances in protein language models (PLMs) have revolutionized IDP research by capturing evolutionary and biophysical signatures at residue-level resolution ^30,42,43,^. While PLMs like ESM-1b/2 and ProtT5 excel in sequence-based functional motif prediction ^31-33,42,43^, AlphaFold2 (AF2) provides structural insights through atomic-level modeling ^27,34,49^. This dual perspective—sequence-derived embeddings versus structure-based features—motivated our systematic comparison to identify optimal representations for multi-category IDR function annotation. Details about PLMs and AF2-derived features, please refer to Supplementary Note 1 and Note 2, respectively.

Benchmarking three PLM-integrated frameworks (IDPFunNet-ESM-1b, -ESM2, -ProtT5) on TE210 revealed architecture-dependent specialization (Figure 3a and 3b). ProtT5 demonstrated cross-category superiority, achieving ≥0.9% higher AUC for protein-binding and ≥4.4% AUC improvement for lipid-binding IDRs, with DFL prediction APS exceeding ESM models by 29.7%. Conversely, ESM2 excelled in nucleic acid-binding task (≥1.1% AUC gain). ProtT5’s average accuracy surpassed ESM-family models by ≥1.5% AUC and 10.2% APS across binding IDR categories, establishing its broad applicability.

**Figure 3.**
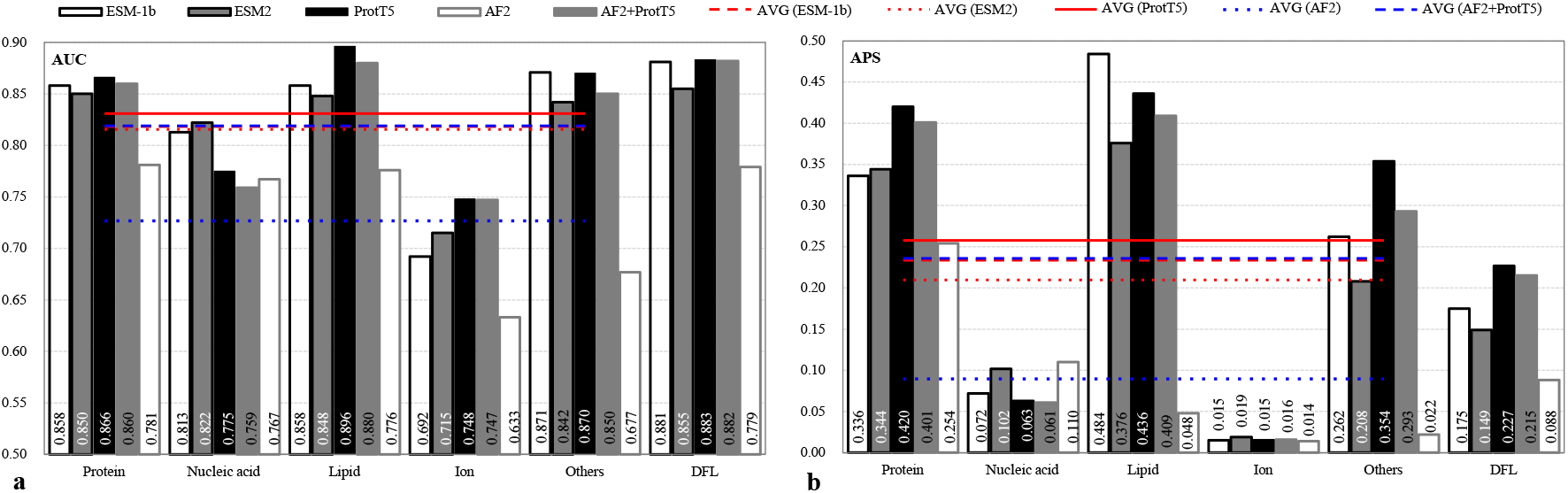
Performance comparison of five IDPFunNet variants on the TE210 benchmark. **(a)** AUC and **(b)** APS metrics demonstrate differential predictive capabilities across six functional categories of IDRs. The “Others” category represents IDRs binding to other small molecules. The cross-category average AVG (*) aggregates performance across five binding classes: protein-, nucleic acid-, lipid-, ion-, and other small molecule-binding IDRs. Model variants employ distinct sequence encoding strategies: three implementations using pretrained language model embeddings (ESM-1b/ESM2/ProtT5) versus two structure-based approaches using AF2-derived features either independently (i.e., AF2) or concatenated with ProtT5 embeddings (i.e., AF2+ProtT5).

Comparative analysis with AF2-derived features highlighted ProtT5’s dominance in five of six functional categories (Figure 3a and 3b). ProtT5 achieved significantly higher AUC in protein-binding (0.866 vs. 0.781), lipid-binding (0.896 vs. 0.776), and DFL prediction (0.883 vs. 0.779), with APS values 15-fold greater in other small molecule-binding. Although AF2 showed marginal advantages in nucleic acid-binding (AUC 0.775 vs. 0.767) and ion-binding (APS 0.015 vs. 0.014), the ProtT5+AF2 hybrid model consistently underperforms ProtT5 alone with relatively small margin (ΔAUC ≤0.02), suggesting limited complementarity between sequence and structural embeddings.

In summary, ProtT5 demonstrates robust cross-category performance (average AUC improvement: 1.2– 14.2%), with its encoder-decoder architecture proving particularly effective in capturing conformational plasticity essential for IDR multifunctional annotation. This enhanced capacity to represent structurally dynamic regions establishes ProtT5 as the optimal choice for IDPFunNet’s sequence encoding framework, where evolutionary-biophysical feature integration drives prediction accuracy.

### Multi-task learning enhances prediction robustness for binding IDR prediction

Existing approaches for IDR function prediction predominantly employ single-task architectures optimized for individual binding types ^25^. This specialization limits their capacity to exploit shared functional patterns across interaction categories.

Our comparison of single-versus multi-task architectures across five functional categories (protein/nucleic acid/lipid/ion/other small molecule-binding) reveals critical performance tradeoffs; Please refer to Figure 4. The multi-task model demonstrated superior AUC in four categories: protein-binding (3.1% improvement), lipid-binding (11.9% increase), ion-binding (28.1% gain), and other small molecule-binding (35.1% higher). Conversely, the single-task architecture excelled in nucleic acid-binding (AUC: 0.817 vs. 0.775; APS: 0.184 vs. 0.063). APS analysis revealed multi-task dominance in protein-binding (0.42 vs. 0.356) and other small molecule-binding (0.354 vs. 0.020), while the single-task model achieved a 0.121 higher APS for nucleic acid-binding.

**Figure 4.**
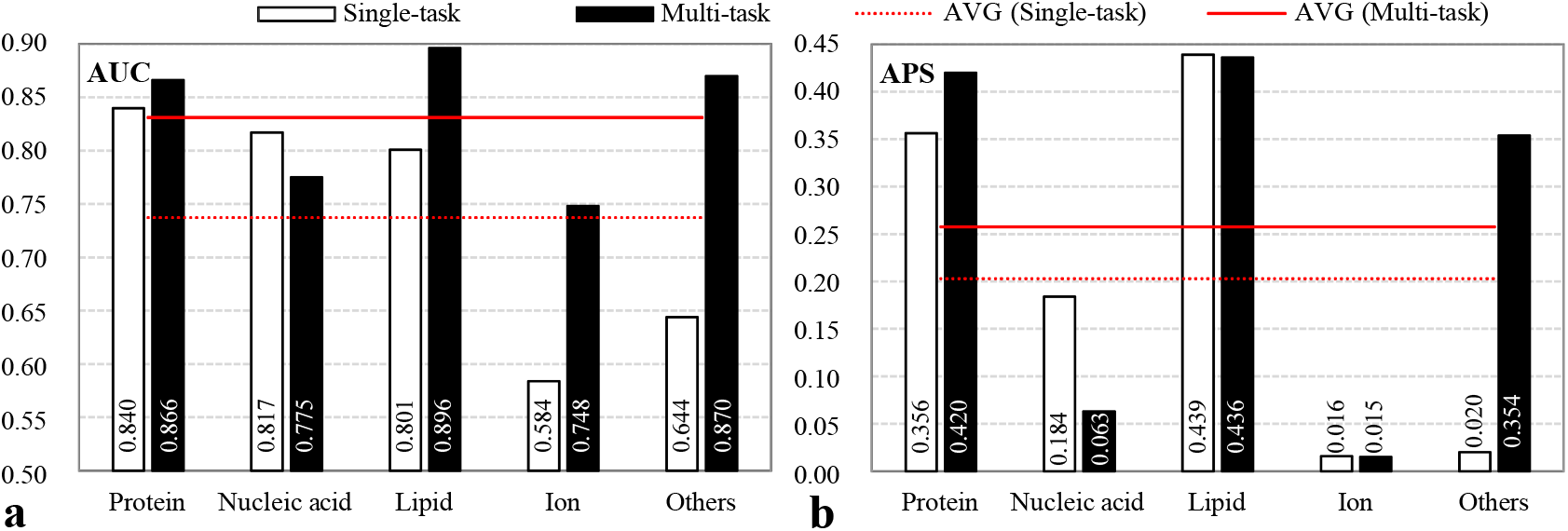
Performance of single-task and multi-task models across five binding types of IDRs, which is evaluated by using (**a**) AUC and (**b**) APS metrics on the test set TE210. The “Others” category represents IDRs binding to other small molecules. AVG (*) aggregates performance across five binding classes: protein-, nucleic acid-, lipid-, ion-, and other small molecule-binding IDRs.

Aggregated across all categories, multi-task learning achieved a 12.7% higher mean AUC and 26.9% greater mean APS compared to the single-task framework. These improvements reflect enhanced feature sharing across correlated tasks, particularly evident in other small molecule-binding where cross-task correlations are strongest.

Therefore, IDPFunNet adopts the multi-task paradigm to balance comprehensive functional coverage with biological fidelity.

### Performance of attention mechanism for IDR multifunction prediction

The multi-head attention (MHA) mechanism uniquely captures long-range sequential dependencies while maintaining local patterns through positional encoding ^50^. Leveraging its demonstrated success in protein structure prediction via transformer frameworks ^34^, we evaluated MHA’s capacity for IDR functional analysis through two engineered variants: 1) MHAmod replaces IDPFunNet’s dual-path architecture with pure MHA modules, and 2) MHAmulti substitutes the dual-path system with a hybrid of multi-scale CNNs (kernel sizes: 1, 3, 5, 7) and MHA — a design proven to extract hierarchical features ^24^. These comparative models were benchmarked against the original IDPFunNet across six functional categories using the TE210 test set, with performance quantified by AUC and APS metrics (Figure 5). Comparative evaluation, shown in Figure 5, revealed distinct performance patterns across functional categories. For binding IDRs, IDPFunNet and MHAmulti achieved superior mean performance relative to MHAmod, with average gains of 0.07 AUC (0.03 APS). This advantage extended to DFL prediction, where both models demonstrated substantial improvements of 0.273 AUC (0.193 APS). Detailed category-specific analysis showed consistent dominance in protein-(AUC: 0.87 vs. 0.77; APS: 0.42 vs. 0.30), lipid-(AUC: 0.89 vs. 0.84; APS: 0.436–0.455 vs. 0.418), and other small molecule-binding IDRs (AUC: 0.87 vs. 0.77; APS: 0.35 vs. 0.22). Notably, MHAmod exhibited exclusive superiority in nucleic acid-binding IDRs (AUC: 0.831 vs. 0.78; APS: 0.195 vs. 0.06), while ion-binding predictions remained challenging for all models (APS<0.02). Despite near-equivalent performance between IDPFunNet and MHAmulti (ΔAUC < 0.01, ΔAPS < 0.02), the former’s streamlined architecture with reduced parameter complexity offers practical implementation benefits.

**Figure 5.**
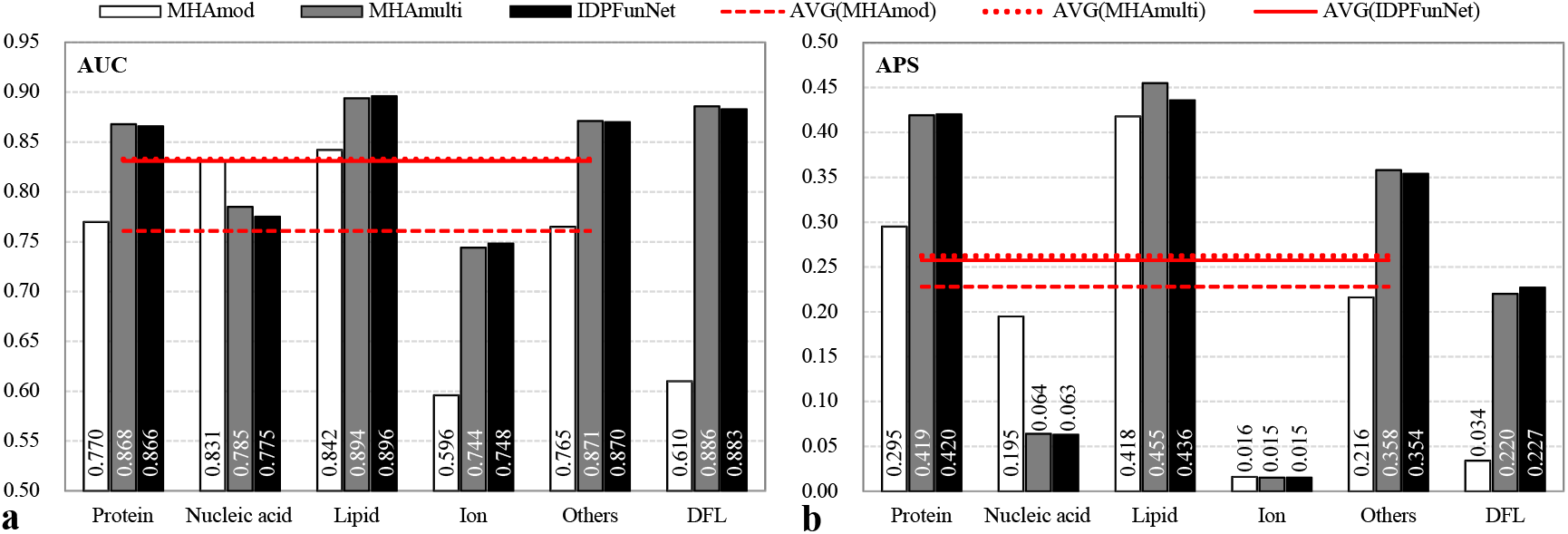
Performance of three models MHAmod, MHAmulti and IDPFunNet across six functional categories of IDRs, which is evaluated by using (**a**) AUC and (**b**) APS metrics on the test set TE210. The “Others” category represents IDRs binding to other small molecules. AVG (*) aggregates performance across five binding classes: protein-, nucleic acid-, lipid-, ion-, and other small molecule-binding IDRs.

In brief, our systematic comparison identifies IDPFunNet as the optimal framework balancing predictive power and architectural efficiency.

### Benchmarking against SOTA methods across IDR functional categories

We evaluated IDPFunNet’s predictive accuracy for DFLs and five binding IDR types across two independent test sets (TE210/TE83), benchmarking against three SOTA predictors: DisoFLAG ^43^, DeepDISOBind ^24^ and DisoLipPred ^25^. DisoFLAG was stated to outperform leading methods in CAID2 ^43^, and was reported as the top-ier in the latest CAID3 (https://caid.idpcentral.org/challenge/results). It employs a protein language model and graph-based interactive architecture to systematically predict DFLs and IDRs interacting with proteins, DNA, RNA, ions, and lipids. DeepDISOBind, a deep convolutional network, concurrently predicts IDRs binding to RNA, DNA, and proteins. DisoLipPred, a bidirectional recurrent neural network, specializes in lipid-binding IDR prediction. Since DisoFLAG and DeepDISOBind are designed to predict DNA-binding and RNA-binding IDRs separately, we combined their prediction scores and binary classifications as the results for nucleic acid binding for a unified evaluation. Please refer to Supplementary Note 3 for details.

IDPFunNet demonstrates robust performance across IDR functional categories, as evidenced by the computational analysis from the TE210 and TE83 datasets (Figure 6, and Supplementary Figure 1-2 and Table 2). On TE210, IDPFunNet attained AUC, APS, and F1-max ranges of 0.748–0.896, 0.015–0.436, and 0.035–0.477, respectively. On TE83, these metrics range from 0.743–0.848 (AUC), 0.037–0.427 (APS), and 0.107–0.450 (F1-max). Holistic evaluation across AUC, APS, F1-max, and MCC revealed that IDPFunNet achieved the highest accuracy for protein binding (AUC ≥0.832, APS ≥0.42, F1-max ≥0.45, MCC ≥0.356), followed by lipid binding (AUC≥0.848, APS ≥0.213, F1-max ≥0.351, MCC ≥0.329), while performance was suboptimal for nucleic acid and ion binding (AUC ≤0.774, APS ≤0.170, F1-max ≤0.276, MCC ≤0.244).

**Figure 6.**
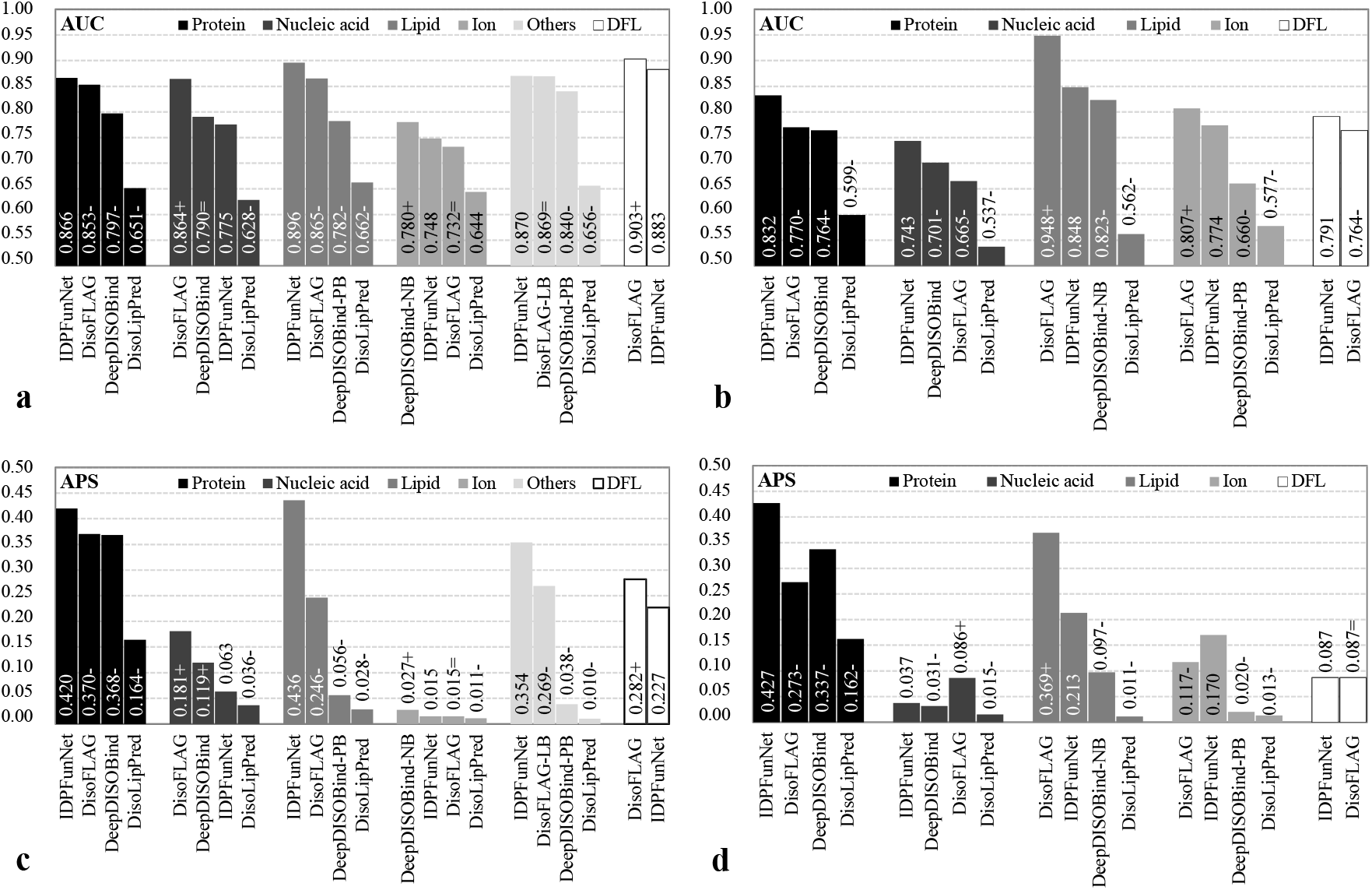
Comparative performance of IDPFunNet and SOTA methods on independent test sets TE210 (**a** and **c**) and TE83 (**b** and **d**), as measured by AUC (**a** and **b**) and APS (**c** and **d**). For lipid-, ion-, and other small molecule-binding IDR predictions, we evaluated model variants DeepDISOBind-PB, DeepDISOBind-NB and DisoFLAG-LB, where PB/NB/LB denote protein/nucleic acid/lipid binding. These variants correspond to task-specific components of DeepDISOBind and DisoFLAG frameworks. Notably, other small molecule-binding IDR annotations are unavailable in TE83. “+”/”−” next to a given AUC and APS value indicates that the corresponding method is significantly better/worse than IDPFunNet (p-value < 0.05).

Comparative analysis with SOTA methods revealed IDPFunNet’s task-specific superiority, achieving dominant performance in protein-binding prediction while maintaining competitive accuracy for other binding types and DFLs. As detailed in Figure 6, and Supplementary Figures 1-2 and Table 2, IDPFunNet attained peak metrics for protein-binding IDRs across both TE210 and TE83 datasets, outperforming SOTA methods by significant margins (improvements: 1.5% AUC, 13.5% APS, 4.8% F1-max, 8.9% MCC). This advantage extended to lipid-binding prediction on TE210, where IDPFunNet achieved 0.896 AUC/0.436 APS/0.453 F1-max/0.429 MCC - surpassing DisoFLAG, DeepDISOBind-PB, and DisoLipPred by ≥3.6% AUC, ≥77.2% APS, ≥48.5% F1-max, and 175% MCC. Nucleic acid-binding performance diverged, with IDPFunNet (AUC: 0.775) trailing DisoFLAG (AUC: 0.864) and DeepDISOBind (AUC: 0.79) on TE210 but showing at least 6% AUC gain on TE83. Notably, IDPFunNet achieved cross-dataset enhancements on TE83: 6% AUC gain for nucleic acid-binding, 43.5% APS improvement for ion-binding, and 3.5% AUC increase for DFLs, alongside 0.198-2.128 times MCC boosts across five binding classes (p-value<0.05).

Moreover, IDPFunNet shows good robustness in our multi-class classification tasks across the TE210 and TE83 datasets (the latter TE83 being a recently updated functional IDR dataset). As illustrated in Figure6 and Supplementary Table 1, IDPFunNet and DeepDISOBind exhibited superior stability in protein binding prediction, with all four metrics’ variations ≤0.05 across datasets. For nucleic acid binding, IDPFunNet showed smaller performance fluctuations (AUC variation <0.05) compared to DeepDISOBind and DisoFLAG, the latter displaying the largest AUC variation (∼0.2). In lipid binding, stability varied by metric: IDPFunNet maintained AUC stability (variation <0.05) but showed APS/F1-max shifts >0.1, while DisoFLAG had minimal MCC variation (0.016) but larger AUC/APS/F1-max changes (0.083–0.138). For DFL prediction, both IDPFunNet and DisoFLAG exhibited significant metric fluctuations (>0.05 variation in all metrics). Collectively, IDPFunNet demonstrated enhanced stability for protein and nucleic acid binding compared to SOTA alternatives.

IDPFunNet establishes superior performance in IDR multifunction prediction, particularly excelling in protein/lipid interaction modeling. Although nucleic acid-binding prediction remains challenging, its cross-task generalizability and dataset adaptability outperform alternatives. The framework demonstrates enhanced robustness for core interaction types despite inherent limitations in data-scarce categories (ion-binding) and class-imbalanced scenarios. These findings position IDPFunNet as a versatile solution for IDR multifunctional annotation, balancing specialized accuracy with broad applicability.

### Benchmarking against CAID top performers for binding IDR and DFL prediciton

To rigorously evaluate IDPFunNet’s predictive capacity for binding IDRs and DFLs, we established a benchmarking framework using four CAID blind test sets (CAID2/3_Bind and CAID2/3_DFL), comparing against the top five AUC-ranked methods from CAID2/3. Prediction results for reference methods were retrieved from the CAID portal (https://caid.idpcentral.org/challenge/results). For binding IDR assessment, we exclusively utilized IDPFunNet’s protein-binding module due to its demonstrated stability across TE210/TE83 benchmarks. Similarly, only the DFL-specific component was evaluated on CAID2/3_DFL sets. Given DisoFLAG’s inconsistent validation performance, we assessed all its binding IDR modules on CAID2/3_bind but reported only the top three performers (by AUC) in Figure 7 and Supplementary Table 2. For DFL comparisons, we included DisoFLAG’s top-ranked DFL predictor DisoFLAG_IDR in CAID3 alongside its dedicated DFL module (DisoFLAG_DFL) to ensure equitable evaluation on CAID2/3_DFL (Figure 7, Supplementary Figure 3-4 and Table 2).

**Figure 7.**
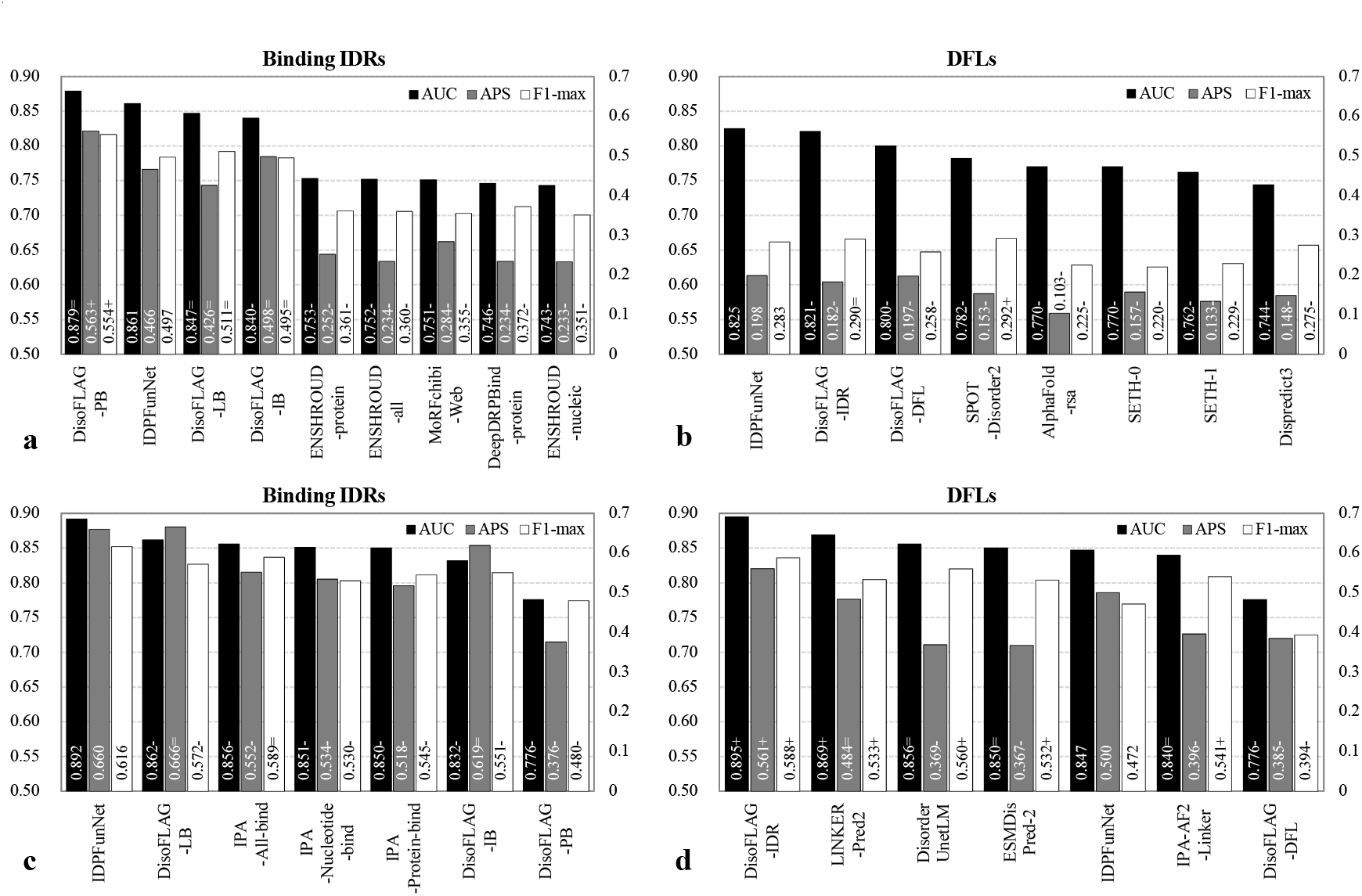
Performance evaluation of IDPFunNet on CAID2 (**a** and **b**) and CAID3 (**c** and **d**) blind test datasets. The *x*-axes present the top 5 AUC-ranked methods in CAID competitions for binding IDR (**a** and **c**) and DFL (**b** and **d**) prediction, alongside DisoFLAG’s top 3 components (**a** and **c**). AUC values are displayed on the primary *y*-axes; APS and F-max on the secondary *y*-axes. The model variants DisoFLAG-LB/IB/PB/DFL denote task-specific components of the DisoFLAG framework, where PB/LB/IB is the abbreviation of protein/lipid/ion binding. “+”/”−” next to a metric value indicates that the corresponding method is significantly better/worse than IDPFunNet (p-value < 0.05). Notably, (**d**) summarizes the results for 14 sequences extracted from CAID3_DFL, i.e., denoted as CAID3_DFL14 set, as all methods listed on the *x*-axis can generate DFL predictions for them.

On CAID2 benchmarks (Figure 7a-b, Supplementary Figure 3 and Table 2), IDPFunNet achieved significant performance gains over all top-five methods (binding: ENSHROUD-protein/all/nucleic, MoRFchibi-Web, DeepDRPBind-protein; DFL: SPOT-Disorder2, AlphaFold-rsa, SETH-0/1, Dispredict3), with minimum AUC/APS improvements of 0.108/0.182. Compared to DisoFLAG, IDPFunNet matched the best-performing DisoFLAG-PB component in binding AUC while surpassing both DisoFLAG-IDR and DisoFLAG-DFL in DFL metrics (0.5% higher AUC and APS).

For CAID3_bind (Figure 7c, Supplementary Figure 4 and Table 2), IDPFunNet outperformed the top five CAID3 methods (DisoFLAG-LB/PB, IPA-All/Nucleotide/Protein-bind) and DisoFLAG’s leading components across almost all metrics (0.03–0.12 higher AUC, -0.01-0.29 higher APS, and 0.03–0.14 higher F1-max). In CAID3_DFL assessment (Supplementary Table 2), IDPFunNet exhibited 0.017/0.061/0.084 improvements in AUC/APS/F1-max over DisoFLAG-DFL, though its AUC trailed top CAID3 methods (IPA-AF2-Linker, DisoFLAG-IDR, etc.) by 0.011–0.042 while exceeding their APS by 0.05–0.169. Notably, critical limitations emerged: leading DFL predictors (IPA-AF2-Linker, ESMDisPred-2) failed to process long sequences (Supplementary Table 2), prompting evaluation on the CAID3_DFL14 subset (14/20 fully predicted proteins). This adjustment altered method rankings (e.g., IPA-AF2-Linker AUC decreased from 0.881 to 0.84), with all reported differences remaining statistically significant (p-value<0.05; Figure 7d, Supplementary Figure 4 and Table 2).

Collectively, IDPFunNet demonstrated robust cross-platform consistency, maintaining top-ier rankings for binding IDR prediction while delivering competitive DFL accuracy (CAID2/3). Intriguingly, generic IDR predictors (DisoFLAG-IDR, SETH series) consistently outperformed DFL-specialized tools (DisoFLAG-DFL, IPA-AF2-Linker), potentially reflecting incomplete DFL annotations in current databases.

## Discussion

IDPFunNet introduces a hybrid deep learning framework for systematic prediction of six IDR functions, encompassing DFLs and five binding subtypes (protein/nucleic acid/lipid/ion/other small molecule). The architecture employs a dual-path design: 1) a CNN-BiLSTM hybrid capturing local motifs and long-range dependencies for multi-class binding predictions, and 2) a dedicated BiLSTM network optimized for DFL annotation. This structural specialization addresses the heterogeneous nature of IDR functions through parallel processing of distinct feature hierarchies.

Benchmarking across six datasets (including CAID2/3 blind tests) demonstrates IDPFunNet’s superior performance in protein IDR prediction, achieving AUCs >0.83 with 13.5–26.7% APS improvements over SOTA methods (p-value<0.05). The model consistently ranked top-two in CAID competitions, validating cross-dataset robustness. For DFL annotation, IDPFunNet maintained competitive accuracy (AUC: 0.791–0.883) across datasets, validating its generalizability.

A key innovation involves ProtT5-based feature engineering, where evolutionary-biophysical residue embeddings outperformed ESM-family models and AlphaFold2 structural features. Integration of ProtT5 enhanced prediction stability (cross-dataset AUC variation ≤0.05) while boosting average AUC/APS by 1.5%/10.2%, respectively. This establishes protein language models as critical for encoding context-dependent binding signatures.

Current limitations include moderate nucleic acid-binding prediction (AUC: 0.743–0.775), where single-task architectures with ESM2 embeddings show superior performance, suggesting subtype-specific optimization needs. Class imbalance in lipid/ion-binding datasets (e.g., TE83: <10 positives) further constrained APS reliability, necessitating synthetic minority oversampling techniques. Future iterations may integrate attention mechanisms for nucleic acid interactions and active learning strategies to address data scarcity.

As a novel framework concurrently addressing multiple IDR functions, IDPFunNet advances disorder biology research through three core strengths: 1) hybrid architecture balancing local/global feature extraction, 2) language model-enhanced evolutionary profiling, and 3) modular design adaptable to emerging functional categories. These innovations position it as a versatile tool for decoding protein disorder-function relationships, with potential applications in pathogenic mutation interpretation and targeted drug discovery.

## Methods

### Model architecture

IDPFunNet represents a novel hybrid deep learning framework designed for comprehensive functional annotation of IDRs, integrating evolutionary sequence information with biophysical properties reflecting structural dynamics to concurrently predict six critical biological functions. As illustrated in Figure 1, the architecture systematically processes input sequences through three hierarchically organized computational stages: sequence encoding, dual-path feature integration, and parallel functional prediction. This design enables simultaneous identification of five distinct interaction types (protein/nucleic acid/lipid/ion/other small molecule-binding) and DFLs, providing a unified platform for probing IDR multifunctionality.

#### Step 1: sequence encoding via ProtT5 embeddings

The framework initiates with ProtT5-based sequence embeddings, where each residue is encoded into a 1024-dimensional feature vector capturing context-sensitive evolutionary patterns and biophysical properties. Selected for its demonstrated capacity to resolve residue-level signatures^33,42,43^, ProtT5 effectively preserves conformational plasticity signatures essential for distinguishing binding motifs from flexible linkers. This encoding establishes the foundational representation for subsequent functional discrimination.

#### Step 2: dual-path architecture for feature integration

The encoded features undergo parallel processing through two specialized pathways optimized for distinct functional categories: (1) A hybrid CNN-BiLSTM path (blue in Figure 1a) processes binding IDR prediction by synergistically combining 2-layer CNN with kernel size 3 for local motif detection ^44^ and bidirectional LSTM networks for long-range dependency modeling ^45^. The CNN component extracts position-invariant binding patterns, while the BiLSTM captures context-dependent sequence correlations, generating a 512-D feature vector from each submodule. (2) A pure BiLSTM path (orange in Figure 1a) dedicated to DFL identification leverages long-range dependency modeling to detect evolutionarily conserved flexible linkers, distinct from binding IDRs ^19^.

#### Step 3: parallel functional prediction blocks

Final predictions are generated through six residue-level classifiers operating on the integrated feature vectors: (1) The binding-specific vector feeds five parallel ResMLP blocks (Figure 1b), each specialized for a particular interaction type (proteins/nucleic acids/lipids/ions/other small molecules); (2) The DFL-specific vector is processed by a dedicated ResMLP block. Each predictor implements a 2-layer ResMLP followed by sigmoid activation, converting feature vectors into position-wise propensity scores:

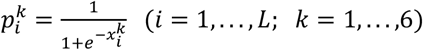

where 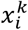 denotes the ResMLP output for residue *i* and function *k*, with 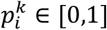 represents functional likelihood. *L* is the length of the input sequence. This parallelized architecture enables computationally efficient multi-task learning while preventing feature interference across distinct functional categories. Details for the deep learning algorithms CNN, BiLSTM, and ResMLP, please refer to Supplementary Note 4.

In summary, IDPFunNet’s architectural innovation lies in its dual-path feature decomposition strategy, which decouples the prediction of binding IDRs from DFLs while maintaining their inherent correlations. By combining global evolutionary patterns from ProtT5 with task-specific local feature extraction, the framework achieves superior performance in capturing IDRs’ context-dependent multifunctionality, as demonstrated in comparative analyses.

### Benchmark datasets

The DisProt database ^56^ serves as a gold standard for IDPs/IDRs, providing extensive and high-quality functional annotations such as binding IDRs and DFLs. These annotations are based on the gene ontology (GO) and the intrinsically disordered proteins ontology (IDPO). Following the GO classifications for protein-, nucleic acid-, ion-, lipid-, and other small molecule-binding, as well as IDPO terms for DFLs, we identified 1,073 sequences with at least one of these six functional annotations from the DisProt database of version 9.4. To ensure annotation accuracy, we refined labels using CAID2’s ^27^ open-source scripts (https://github.com/BioComputingUP/caid2-reference/blob/master/src/references.ipynb), which standardize functional assignments across datasets. Given the difficulty in distinguishing RNA and DNA binding subtypes for nucleic acid binding, and to ensure adequate data for training and testing, nucleic acid binding is not divided into DNA and RNA binding subtypes.

Subsequently, we applied the program CD-HIT ^57^ to cluster the proteins with at least 25% sequence identity. The resulting 814 clusters were randomly divided into training, validation, and testing datasets. This division ensures the independence among these three datasets at the level of 25% sequence identity. Specifically, the training, validation, and test datasets consist of 552, 227, and 210 sequences, respectively, belonging to 463, 198, and 153 clusters. For convenience in subsequent sections, these datasets are denoted as TR552, VA227, and TE210, respectively.

To further investigate the robustness and prediction accuracy of the models, we collected 83 newly deposited sequences from the DisProt database, which were added after version 9.4 but before version 9.6. These sequences form the independent test dataset TE83. Additionally, we included four blind test datasets officially provided by the CAID2 and CAID3 competitions, denoted as CAID2_Bind, CAID2_DFL, CAID3_Bind, and CAID3_DFL. The CAID2_Bind and CAID3_Bind datasets focus on generic binding events mediated by intrinsic disorder, comprising 78 and 51 proteins, respectively, each with at least one IDR binding to protein, nucleic acid, ion, lipid, and/or other small molecules. The CAID2_DFL and CAID3_DFL datasets consist of 40 and 20 sequences, respectively, where IDRs serve as flexible linkers. Notably, each of these four datasets shares less than 25% sequence identity with both our training dataset TR552 and validation dataset VA227.

In summary, the datasets TR552 and VA227 were used for training, while six independent test datasets, including TE210, TE83, CAID2_Bind, CAID2_DFL, CAID3_Bind, and CAID3_DFL, were utilized for model assessment and/or comparison. For more detailed information about these benchmark datasets, please refer to Supplementary Table 3.

### Training process

We separately trained the hybrid CNN-BiLSTM path for binding IDR prediction and the BiLSTM-based path for DFL prediction, to optimize the IDPFunNet model. When training the hybrid CNN-BiLSTM path, we define the loss of the generic disordered binding *Loss*_*DB*_ as below:

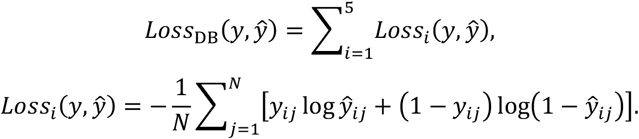

*Loss*_*i*_ (*i* = 1,2, *⋯*, 5) is the binary cross-entropy loss function, where *i* represents a given type of disordered binding event, i.e., the IDRs interacted with protein, nucleic acid, lipid, ion, or other small molecules. We take each input sequence as a batch, and thus *N* corresponds to the total number of residues in the input sequence. *y* is the real labels, while *ŷ* means the predicted scores. For the BiLSTM-based path, we also utilized the binary cross-entropy loss *Loss*_6_ (6 representing to DFLs) by following the formula for *Loss*_*i*_ (*i* = 1,2, *⋯*, 5). We use the pytorch framework to train these two paths with an optimizer of stochastic gradient descent (SGD) and a learning rate of 5e-4. We choose the network when the sum of the losses across all batches (i.e., all training sequences) is minimized.

### Evaluation metrics

IDPFunNet predicts six numeric propensity scores to quantify the likelihood of a residues located in DFLs or the IDRs binding to proteins, nucleic acids, lipids, ions and other small molecules. Following CAID competitions ^26,27^, we evaluate its predictive performance using AUC, APS and F1-max.

We first sort the predicted probabilities of the positive class to obtain a set of probability thresholds 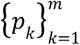 (*m* is the total number of unique thresholds). Then, we calculate true positive rate (*TPR*), false positive rate (*FPR*), precision (*PRE*) and recall (*REC*) at each threshold *p*_*k*_, using standard formulas:

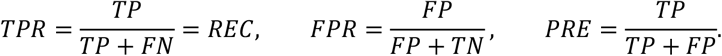

AUC, the area under the receiver operating characteristic (ROC) curve constructed by (*FPR, TPR*) points, assesses overall classification performance. A value close to 1 implies excellent performance, 0.5 indicates random guessing, and less than 0.5 shows sub-par performance.

APS refers to the weighted average of the *PRE* values at different *REC* levels, and is calculated as:

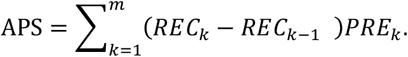

A high APS means the model is good at finding correct instances and minimizing false detections. F1-max means the maximum value of the F1-score across all thresholds *p*_*k*_ (*k* = 1,2, …, *m*). F1-score is a harmonic mean between *PRE* and *REC*, which is formulated as:

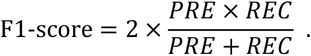

Generally, a higher F1-max indicates better overall classification. Additionally, F1-max often presents a strong positive correlation with APS.

IDPFunNet also offers binary predictions. Using the probability threshold 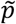-corresponding to F1-max, residues with 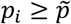-are classified as positives (disordered and functional), and others are negatives. We use MCC to assess the quality of binary predictions^19,21,22,24^. MCC is calculated from all four confusion matrix elements (*TP, TN, FP*, and *FN*) as below:

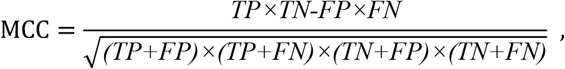

with values ranging from -1 to 1. A MCC of 1 means perfect prediction, 0 implies random guessing, and -1 indicates total misclassification. It is especially reliable for imbalanced datasets, and a higher value indicates better classification.

### Hypothesis testing framework

To rigorously evaluate the statistical significance of IDPFunNet’s performance advantages, we implemented a hypothesis testing framework across six independent test sets using four evaluation metrics: AUC, APS, F1-max, and MCC. Our protocol initiated with bootstrap resampling (30 iterations per test set), where each iteration randomly selected 50% of sequences while preserving the original positive/negative class distribution. For categories with limited positive samples (<10 instances, e.g., TE83 lipid-binding), we retained all positive cases and proportionally subsampled negatives to maintain the 50% sequence threshold. This approach generated paired 30-dimensional metric vectors (IDPFunNet vs. comparator methods) for subsequent statistical analysis.

Prior to hypothesis testing, we assessed data normality using the Anderson–Darling test (α=0.05). Normally distributed vectors underwent paired t-tests, while non-normal distributions were analyzed via Wilcoxon rank-sum tests. Statistical significance was established at p-value<0.05, ensuring robust validation of IDPFunNet’s superior predictive capabilities across both DFLs and specific/generic binding IDRs.

Notably, technical constraints affected comparative analyses for certain methods (e.g., IPA-AF2-Linker, ESMDisPred-2), which failed to generate predictions for 30% (6/20) of CAID3_DFL sequences and are currently inaccessible. To maintain evaluation fairness, we restricted hypothesis testing to the 14 CAID3_DFL proteins with complete predictions across all benchmarked methods. This adjustment eliminated potential bias from missing data while preserving statistical power through consistent sample availability.

## Code availability

IDPFunNet is freely available at https://github.com/IDRIDP/IDPFunNet/tree/master and https://yanglab.qd.sdu.edu.cn/IDPFunNet1/.

## Acknowledgments

This work was supported in part by National Key Research and Development Program [NK R&D Program 2023YFF1204003] and National Natural Science Foundation of China grants [NSFC T2222012]

